# Murburn concept explains the acutely lethal effect of cyanide

**DOI:** 10.1101/555888

**Authors:** Kelath Murali Manoj, Surjith Ramasamy, Abhinav Parashar, Vidhu Soman, Kannan Pakshirajan

## Abstract

Cyanide (CN) toxicity is traditionally understood to result from its binding of hemeFe centers, thereby disrupting mitochondrial cytochrome oxidase function and oxygen utilization by other globin proteins. Recently, a diffusible reactive oxygen species (DROS) mediated reaction mechanism called murburn concept was proposed to explain mitochondrial ATP-synthesis and heat generation. Per this purview, it was theorized that CN ion-radical equilibrium dissipates the catalytically vital DROS into futile cycles, producing water. In the current study, a comparative quantitative assessment of the above two explanations is made for: (i) lethal dosage or concentrations of CN, (ii) thermodynamics and kinetics of the binding/reaction, and (iii) correlation of CN with the binding data and reaction chemistry of H_2_S/CO. The quantitative findings suggest that the hemeFe binding-based toxicity explanation is untenable. CN also inhibited the experimental *in vitro* DROS-mediated coupling of inorganic phosphate with ADP. Further, pH-dependent inhibition profiles of heme enzyme catalyzed oxidation of a phenolic (wherein an -OH group reacts with DROS to form water, quite akin to the murburn model of ATP synthesis) indicated that- (i) multiple competitive reactions in milieu controlled outcomes and (ii) low concentrations of CN cannot disrupt activity via a coordination (binding) of cyanide at the distal hemeFe. Therefore, the μM-level IC_50_ and the acutely lethal effect of CN on cellular respiration could be explained by the deleterious interaction of CN ion-radical equilibrium with DROS in matrix, disrupting mitochondrial ATP synthesis. This work supports the murburn explanation for cellular respiration.

## Introduction

Since three centuries or so, cyanide has captivated human attention post the synthesis/discovery of the dye Prussian blue (ferric ferrocyanide, Fe_7_[CN]_18_) and prussic acid (hydrogen cyanide, HCN). Owing to its lethality, cyanide salts and acid (hereon termed CN) have been regulated by governments. Though CN-toxicity is well studied, the reason for its acutely lethal effect on cell/organism mortality is obscure. The classical perception on CN-toxicity entails a stoichiometric binding-based mechanism blocking functional heme centers, thereby preventing oxygen utilization by vital cellular heme proteins.^1–2^ Recently, a radical perspective for oxygen utilization, murburn (abstracted from ‘mured burning’) concept, was proposed to explain the physiology of xenobiotic metabolism, maverick physiological dose responses and cellular respiration.^3–10^ The new murburn proposal states that diffusible reactive oxygen species (DROS) generated at membrane interfaces are crucial electron transfer and catalysis agents in routine physiology. In cellular respiration or oxidative phosphorylation (OxPhos), the murburn explanation vouches that DROS generated at mitochondrial complexes directly serve as the coupling agents for ADP and Pi, leading to ATP synthesis. In this scheme, it was proposed that cyanide ion-radical mediated catalysis, and not cyanide binding to heme (as currently held), is the primary rationale for cyanide lethality. This article critically explores the theoretical and experimental premises of the two molecular explanations (as shown in Figure 1) for cyanide-induced acute toxicity. The variables and the quantitative aspects of stoichiometric, thermodynamic, and kinetic parameters for binding-based explanation of various molecules/ions with heme proteins are investigated. Ratifying the predictions of murburn explanation for OxPhos,^11^ we also provide direct experimental evidence for DROS-assisted ATP-synthesis and the inhibition of this process by CN. The murburn explanation is further corroborated with global analyses of select heme-enzyme K_d_-IC_50_ experimental data/profiles.

**Figure 1.**
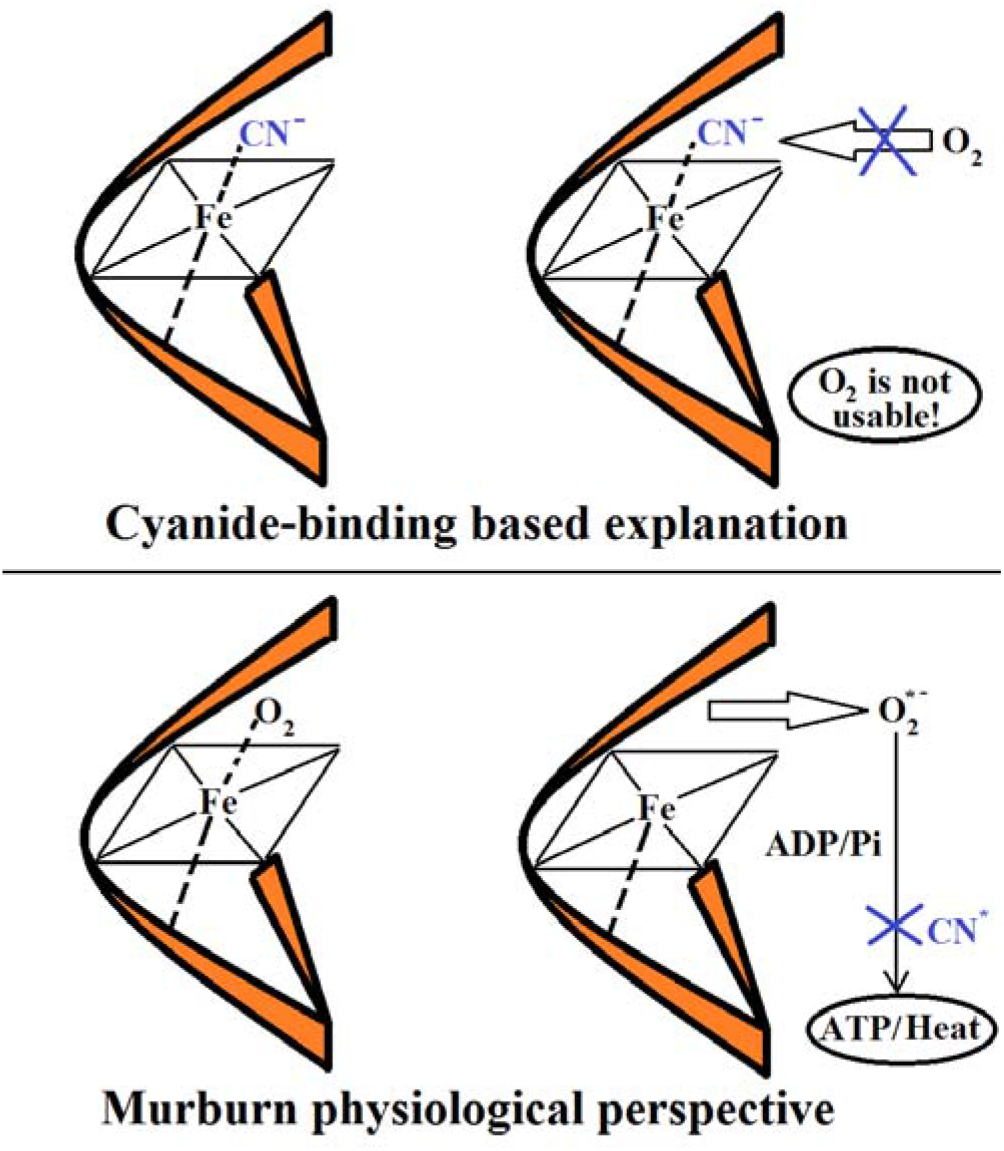
A scheme of the two mechanistic explanations for the toxicity of cyanide: In the textbook explanation, CN ligates to heme-center, thereby preventing oxygen’s accessibility, transport or reduction thereafter. This is a 1:1 reaction outcome, resulting out of a chemico-physical binding. In the murburn purview, cyanide (radical) reacts with the catalytic DROS generated/stabilized at heme-center, thereby disrupting ATP-synthesis. This is a mechanism wherein cyanide acts as a recyclable catalyst.

## Results & Analyses

### Dosage calculations based on stoichiometric CN binding

Following is the dosage analysis of cyanide, based on human blood’s oxygen-transporting capacity (A) and Cox’s oxygen-binding ability (B).

**A**. *Blood’s oxygen transporting capacity.* Each RBC contains ~10^9^ CN-binding sites (= 2.5 × 10^8^ Hb × 4 sites per Hb) and since one liter of blood contains ~5×10^12^ RBCs, 5 liters of blood in a human has ~ 2.5 × 10^22^ binding sites of CN. As blood levels of ~3 mg/L cyanide is accepted to be lethal, the threshold blood load is 15 mg of cyanide.^12^ Since 26 g (ionic mass) of cyanide contains 6.023 × 10^23^ cyanide ions, 15 mg contains [(6.023 × 10^23^) × (15/26)/1000] = 3.5 × 10^20^ cyanide ions. Granting the best-case scenario of 100 % irreversible binding efficiency, the CN lethal load is only a minute fraction (1.4 %) of the total binding sites in blood. This leads to the inference that cyanide-lethality is not brought about by the disruption of Hb-O_2_ interactions. A similar analysis also downplays CN-myoglobin binding-based outcomes. Also, since 4-dimethyl aminophenol (DMAP) is used as an antidote for CN-poisoning^13^ and it is known to work by generating met-species of the blood/muscle globins (which in turn facilitates CN-binding by these proteins!), acute toxicity of low doses of CN via inhibition of oxygen transport or storage is ruled out.

**B**. *CN-binding to the heme-center of Cox*: An average human could have 20 to 70 trillion cells. Considering ~10^7^ Cox per cell (= 10^4^ Complex IV per mitochondrion × ~10^3^ mitochondria per cell),^14^ the estimate for the total Cox binding sites is >4 × 10^20^. Across several organisms/animals (including humans), KCN is lethal when administered (orally or intra-venously/peritonially/muscularly) at ~1 mg/Kg.^15–16^ Assuming complete absorption and equi-distribution within a body mass (of equal density as water), the toxic physiological concentration of CN comes to ≤ 10^−6^ M (mg/Kg or mg/L). *In vivo* or *in situ*, the actual and effective concentration of CN would be far lower than what is administered, owing to loss through non-absorption (in the gut) and metabolism/excretion (by say, kidneys). This implies that the ~10^20^ CN present in the mg/Kg administration (calculated in point A above) would be inadequate for lethality even if the binding to Cox is 100 % irreversible and efficient.

### CN binding in physiological regimes and the competitive scenario therein

The fact is that cyanide-binding to heme-centers is neither irreversible nor efficient in physiological realms. The thermodynamic equilibrium dissociation constant K_d_ is defined as the concentration of CN at which half of the total available heme-centers is populated by the ligand (CN). Usually, when micromolar levels of protein is taken, at least 10X K_d_ concentration of ligand is minimally required to saturate >90% of the total active sites. Even in such conditions, binding is via a non-covalent and coordination scheme of ‘off-now, on-now’ kind of interaction, and it is not an ‘always bound’ scenario.^17^ As measured by non-invasive spectroscopic assays in pure solutions, cyanide binding to micromolar levels of hemeproteins show K_d_ values ~10^−3^ M.^18–20^ In particular, the affinity-based K_d_ of cyanide for Cox approaches the mM ranges (similar to that of oxygen) and cyanide has relatively low affinities for the oxidized form of Cox.^20^ Furthermore, binding of small species to membrane-bound proteins with deep-seated active sites would be relatively slow, owing to constraints imposed by diffusion.^19^ Such a reality translates to the conservative requirement of g/Kg or g/L dosage administration for achieving Cox hemeFe binding-based lethal effects, thereby confirming the calculations in point B above. Thus, the stoichiometric binding-based calculations fall short of explaining the theoretically required dosage outcomes by several orders of magnitude. This inference is consolidated by the facts/reasoning that-(i) Not all of the cyanide presented is assimilated by the body and therefore, the calculation is a conservative approximate, with benefit of doubt given to the binding-based explanation, (ii) If a small amount of cyanide is injected (say, in the toe), then it should only render the foot as “reversibly dead”. However, the animal dies quickly and this process is acute and irreversible, (iii) CN is acutely lethal when presented at very low dosages via various modalities (breathing, oral, contact, injection, etc.) to the vast majority of life forms of diverse masses and volume/surface area ratios, (iv) The lethal doses apply even for unicellular organisms and cellular suspensions, (v) CN inhibits certain heme enzymes’ activities even in simple *in vitro* setups at micromolar levels, which cannot be explained by the higher K_d_ values of the protein-ligand interaction.^18^ (vi) The ionic species of CN^−^ would not be readily available for the membrane-embedded Cox.

Physiologically, at the lethal cyanide concentration of ~ 10^−6^ M, the symmetric diatomic oxygen molecule should out-compete the asymmetric triatomic HCN or diatomic ionic cyanide for binding Cox. This is because the hydrophobic oxygen is present at >10^−4^ M in the aqueous milieu and oxygen is almost five times more soluble in the membrane milieu than in water. Experimentally, when 1 μM Cox is taken along with 10^−4^ M cyanide, and then aerated buffer is mixed in a stopped flow apparatus, Cox heme might not show any spectral change within seconds.^21^ This is because at these “unrealistically high cyanide” ratios, Fe-CN binding may prevail! Such an observation cannot ratify the physiological competing ability of low doses of CN vis a vis oxygen. When high levels of the efficient ligand CO bound to blood haemoglobin is easily reversed (by aerating the intoxicated patients), the competitive binding efficiency of CN would only be lower. The ligation of cyanide to hemeFe results via the carbon atom of the CN^−^ anion. Since the pK_a_ of HCN is 9.4 (two logarithmic units higher than the physiological pH), only a minute fraction is available as cyanide ion. This implies that only a small fraction of the low amounts of CN presented would bind to Fe-centers at physiological regimes. Therefore, disruption of heme enzymes by binding and transport based phenomena appear to be unlikely explanations for CN toxicity.

### Comparison of equilibrium and kinetic constants of CN with other small toxic heme ligands like CO and H_2_S

The heme-ligand interactions for Hb or Cox interaction with oxygen can be theorized as a minimal scheme of two steps 1 and 2, as given in Box 1. The original heme concentration is [Heme], the formed complex is [Heme-L] and the free ligand in milieu is [L], Then, for [Heme] < [L] (that is, the ligand in milieu is in far excess of the hemeprotein) and assuming a 1:1 stoichiometry of Heme:L binding (a single ligand binding site per hemeprotein), the laws of equilibrium and steady-state approximations afford the correlations (the famous Hill / Michaelis-Menten / Cheng-Prusoff equations) given in Box 1. The italicized small letter ***k*** is the pertinent rate constant (with units of s^−1^ or M^−1^ S^−1^, depending on whether the reaction is first order or second order) whereas the capital K is the relevant equilibrium constant (in M). The forward reaction rate constants are *k*_1_ and *k*_2_ whereas the backward rate constants are *k*_-1_ and *k*_-2_ (the latter, *k*_-2_, is deemed irrelevant for the current discussion now). The first step is the equilibrium interaction characterizing the K_d_ (dissociation constant) or K_s_ (association constant). The second step is a chemical reaction, which plays a significant factor in the determination of the value of the functional interaction constant K_M_.

Now, from literature, it is evident that the functional K_M_ of oxygen for cytochrome oxidase is < 0.1 μM, which is several orders lower than the experimental K_d_ of ~0.3 mM!^22–24^ From Box 1, it can be seen that this is theoretically non-permissible (as K_M_ = K_d_ + /k_2_/k_1_). That is, K_M_ can only be equal to or greater than K_d_. This simple quantitative analysis successfully challenges the supposition that “oxygen remains bound at the heme-active site of Cox to make water” by the ‘proof by contradiction’ approach. As a consequence, CN inhibition cannot be explained based on a “quantitatively unsound kinetic theory”. It is relevant to mention here that in the various hemeperoxidases and mixed oxidase (cytochrome P450) systems, similar prevailing fallacies were pointed out and the kinetic/mechanistic phenomena were explained by murburn concept.^3–5,10,18,19,25^ Now, overlooking the “non-applicability of binding-based” kinetics, we can analyze the affinity based equilibrium constants for Cox hemeFe. Table 1 is a compilation of original experimental data from various reliable sources, along with some calculated K_d_ values (K_d_ = *k*_-1_/*k*_1_) for corroboration.^20,26–31^ The practical determination of the backward reaction rate constant *k*_-1_ is fraught with errors (particularly more so for Cox!) and therefore, it is not given high importance here.

**Table 1:** A compilation of equilibrium, inhibition and the forward binding rate constants for Cox at physiological pH.^20,26–31^

| Constants | O<sub>2</sub> | CO<br>(Comp.) | HCN<br>(Non-comp.) | H<sub>2</sub>S<br>(Non-comp.) |
| --- | --- | --- | --- | --- |
| Exp. $K_d$ [M] | $2.8 \times 10^{-4}$ | $4 \times 10^{-7}$ | $5.5 \times 10^{-4}$ | $4 \times 10^{-5}$ |
| $K_d = (k_{-1} / k_1)$ [M] | $(2.1 \times 10^{-4})$ | $(1.7 \times 10^{-7})$ | $(5 \times 10^{-7})$ | $(4 \times 10^{-8})$ |
| $K_M$ or $K_i$ [M] | $<1 \times 10^{-7}$ | $3 \times 10^{-7}$ | $2 \times 10^{-7}$ | $2 \times 10^{-7}$ |
| IC<sub>50</sub> [M] | NA | $4 \times 10^{-5}$ | $5 \times 10^{-5}$ | $5.3 \times 10^{-5}$ |
| $k_1$ [M<sup>-1</sup> s<sup>-1</sup>] | $1 \times 10^8$ | $8 \times 10^4$ | $1.3 \times 10^2$ | $1.5 \times 10^4$ |

It is known that ferro-porphyrin hemes demonstrate similar binding profiles for diatomic ligands that induce a strong field splitting of the Fe atom’s *d*-orbitals, like CO and CN. Both CN and CO are sigma-donors and pi-acceptors. The way a ligand binds, i.e. front-on or side-on, also depends on the steric hindrances in the active site and on the molecular orbitals’ overlaps. The diatomic gaseous molecule of CO is isoelectronic to oxygen but is more soluble in the phases involved (and therefore, is more readily available) and is known to be a much stronger ligand of ferrohemes (than oxygen). The longer Fe-C lengths in Fe-CN complexes indicate that CN is a poorer pi-acceptor than CO (although the former may be a better sigma-donor, which could make CN stabilize higher oxidation and low spin states of the metal atom). Thus, it is an accepted scientific consensus that the Fe-CN ligation is weaker than Fe-CO bonding, for most hemeFe systems. In fact, CO can effectively replace CN in several ferro-porphyrin complexes.^32^ The triatomic gas H_2_S is symmetric, smaller and more hydrophobic than HCN, making it readily accessible across biological lipid membranes. Also, H_2_S has a pK_a_ of 7.04, whereby significant amounts of SH^−^ is also present in aqueous physiological milieu, which must make it a more potent inhibitor. Now, glancing across the experimental equilibrium binding constants of the four gases for Cox (the first data row of Table 1), it is noted that only CO has a markedly higher association with the respiratory protein. The other three gases have comparable affinities. When the equilibrium binding K_d_ is extrapolated for the toxic gases from the kinetic constants listed by Cooper & Brown,^17^ H_2_S is marked out as the major inhibitor. This is an inference which has very little physiological significance because it is a known fact that cyanide is a far more potent toxic principle than CO or H_2_S. (The error-prone determination of *k*_-1_ is evident in Table 1, as the calculated K_d_ for the non-competitive inhibitors do not agree with the experimental K_d_.) Further, H_2_S & CO pose identical Ki and IC_50_ to that of CN. The natural questions to be asked are-How can the binding-based explanations help us understand the toxicities? If the binding/interaction constants are comparable, then why is cyanide immensely more toxic? If we take the forward kinetics-based data alone independently, then also H_2_S and CO should be more potent toxic principles for aerobic organisms, which we know is not the case. Most animals can take in several breathes of H_2_S and CO without getting significantly affected whereas a few whiffs of HCN would incapacitate them. Exposure to CO is common place in modern life, and the physiologic background is ~1% CO-Hb in normal people (per point A, this approaches the equivalent load of cyanide needed to kill a human!). Even ~ 10 % CO-Hb does not cause any appreciable short term adverse effects in humans. Only ~50 % CO-Hb leads to a loss of consciousness and >80 % CO-Hb is acutely fatal.^33^ Now, the fallacy of the binding-based explanation is clearly evident. Since CO (or H_2_S) is a better binder, it should be more toxic, which is not the physiological case! Therefore, by comparing with the gaseous ligands of CO and H_2_S, it is beyond reasonable doubt that the old binding-based hypothesis is unviable for explaining the disruptive efficacy of cyanide. Other traditional explanations that lead to specific organ or tissue failures and death thereafter stem from the molecular mechanisms related to events that follow hemeFe-CN binding. Therefore, it is concluded that hemeFe binding-based outcomes cannot explain the acute physiological toxicity of CN.

### Superoxide aided ATP synthesis and inhibition of this process by cyanide

Using the indirect chemiluminescence method of ATP detection (with luciferase enzyme), we had recently reported the DROS-assisted *in vitro* synthesis of ATP (starting from ADP and Pi).^8^ Herein, we traced the *in vitro* DROS-assisted ATP synthesis with a more direct method (and with elaborate controls), using reverse-phase HPLC. Figure 1 of Supplementary Information shows the sample chromatograms of controls (known ADP + ATP mixture) and sample test reactions (ADP + Pi + DROS ± CN). In the controls and tests, clear separation of ADP and ATP peaks is seen. We have thus confirmed the DROS-assisted ATP-synthesis in aqueous milieu and the ATP formation levels agreed with our earlier reported data.^8^ Further, in the test reactions, the disappearance of ADP corresponded to the formation of ATP. Very crucially, the incorporation of millimolar levels of cyanide inhibited even the *in vitro* DROS-assisted ATP synthesis.

**Table 2:** Estimation of ADP and ATP in reaction milieu by HPLC

| No. | Description of sample | Estimated values |  |
| --- | --- | --- | --- |
| | | ADP (μM) | ATP (μM) |
| 1 | Method check: Positive control (200 μM ADP + 100 μM ATP) | 213* | 109* |
| 2 | Negative control: 10 mM Pi ± KO<sub>2</sub> | 0 | 0 |
| 3 | Negative controls: 200 μM ADP ± (KO<sub>2</sub> or 10 mM Pi) | 211 | 0 |
| 4 | Reaction: Test 1 (200 μM ADP + 10 mM Pi + KO<sub>2</sub>) | 202 | 2.1 |
| 5 | Reaction: Test 2 (200 μM ADP + 10 mM Pi + 10 mM CN + KO<sub>2</sub>) | 207 | 0.7 |

\*A slight overshoot is noted (~6-9 %) in quantitative estimation from the slope value of plot of standards prepared earlier. (This could be owing to a slight evaporation of water from the standards.) Otherwise, estimation showed less than 10% variation in duplicate samples/assays.

### Heme-enzyme inhibition profiles demonstrate the crucial affects/effects of proton-availability on CN-superoxide dynamics

Experimentally, we investigated the outcome of various CN and pH levels in the heme-enzyme reactions, particularly for the one-electron oxidation of the phenolic compound, pyrogallol. [[Both HRP (horseradish peroxidase) and catalase show significant oxidase activity. In the murburn reaction scheme for the model system, pyrogallol’s –OH group would be attacked by the DROS as the first step (finally leading to purpurogallin formation). This is quite analogous to the mitochondrial OxPhos where DROS generated by the membrane proteins attack ADP’s/Pi’s –OH group (finally leading to the esterified product of ATP). Both these reactions also form water. Therefore, this catalase/peroxidase–pyrogallol–peroxide reaction system can be taken as a simple physiological model for the OxPhos reaction. This model obviates the need for membrane embedded complex system such as Cox, whose functional reaction of “proton pumps + water formation” is practically difficult to trace experimentally. Please refer Box B, Supplementary Information, for details of the reaction/models studied.]] At pH 5, the pK_a_ of superoxide (4.8) is approached whereas at pH 9, the milieu is closer to the pK_a_ of HCN (9.4). Though the physiological pH is 7.4, the functional steady-state regime in the matrix is practically proton-deficient. (This steady state scenario is equivalent to a higher pH regime, which is owing to the “closed reactor” effect brought out by the membranes and the uniqueness of NADH, a “2e + 1H” donor.) Figure 2 shows the dose response plots of data obtained at pH 9 (which may be compared with the data points/profiles at pH 5 and pH 7, duly provided in Figure A of Supplementary Information). At pH 5, the drop from ~80-75% to ~20-25% activity (as indicated from the traditional Hill Slope), also the “functionally dynamic” range (where experimental rates are more accurately determined with respect to the controls), is achieved within a single decade of cyanide concentration. In contrast, it takes about three decades of cyanide concentration for the same outcome at pH 9. That is, as the pH increases, the “Hill Slope” flattens. Experimental coordination K_d_ calculated for nM levels of heme-enzyme to bind small ligands like CN should approach high decades of mM levels.^18^ However, the functional K_d_ value calculated/deduced for CN-heme approaches low μM levels at nM levels of CN. For example, in Figure 2, the K_d_ of Catalase-CN interaction is calculated to be 2.5 μM as follows-2 nM catalase and 1 μM CN yield ~72% activity of the control (without CN). So, per the scheme of Box A, Supplementary Information, functional K_d_ at this regime = [free heme] [free CN] / [heme-CN] = (0.72 × 2) (1000 – 0.56) / (0.28 × 2) = 2.5 μM. The highly reproducible data we obtained show that such dynamic K_d_ values change by orders of magnitude (upon changing the CN concentration) at pH 9, for both catalase and peroxidase. The K_d_ values were smaller at low cyanide concentrations (Figure 3 and Table A, Supplementary Information), and this is inconsistent with the theory of binding and the concept of K_d_. Such an anomaly with CN was deduced from the analyses of data for many heme enzymes and substrates^18,19^ and seen even at pH 7 (Table A, Supplementary Information). As HRP is monomeric, cooperativity is ruled out as a cause for the observed effects. Since the allosteric modulation by CN binding at other site(s) on the apoprotein is actually incompatible with the distal hemeFe ligation theory, there is no real theoretically or mechanistically sound explanation in the classical purview. As a consequence, multiple competing reactions in milieu (the murburn explanation) seem as the only appealing option left for explaining the observations/results. Figure 3 shows that the slopes flatten with the increase of pH for the various substrates with HRP. Except for the HRP-ABTS reaction (which needs the direct involvement of excess protons in the reaction scheme, as shown in Box B, Supplementary Information), the IC_50_ value falls at pH 7, with higher values at the two pH extremes. This is explained with the fact that the pK_a_ of the two protagonists (superoxide and cyanide) fall close to these respective pH ranges. Only ABTS needs protons’ direct consumption in the reaction and therefore, shows a high IC_50_ at neutral pH. At alkaline pH, HRP-ABTS catalytic cycle is subject to the proton limitation, owing to effective competition by CN. This inference is corroborated by the finding that at acidic pH (as the regime approaches the pK_a_ of superoxide), once again the IC_50_ values are higher. Clearly, the outcomes result owing to complex equilibrium catalyses involving protons and diverse species present/generated in milieu.

**Figure 2:**
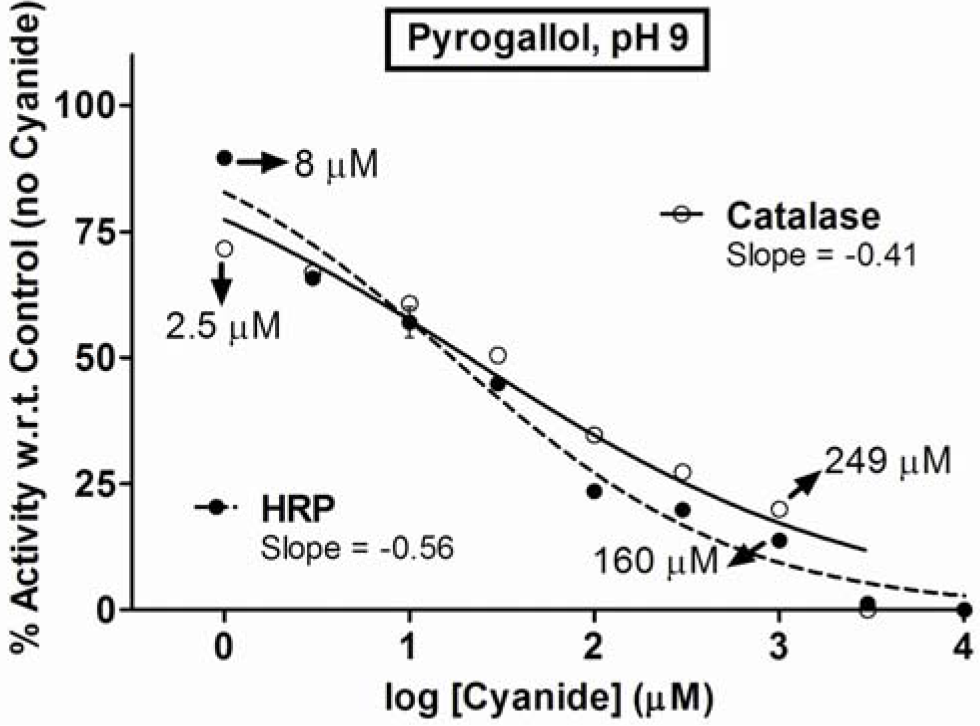
CN sponsored inhibition of the formation of purpurogallin (resulting from the one-electron oxidation of pyrogallol) by horseradish peroxidase (HRP) and catalase at alkaline pH. Functional K_d_ values estimated at the respective four points (marked with arrows) seem to erroneously indicate that “affinity” increases at low cyanide concentrations. (For more details, please refer discussion in the text and Supplementary Information.) Clearly, a slope-flattening is discernible at higher pH.

**Figure 3:**
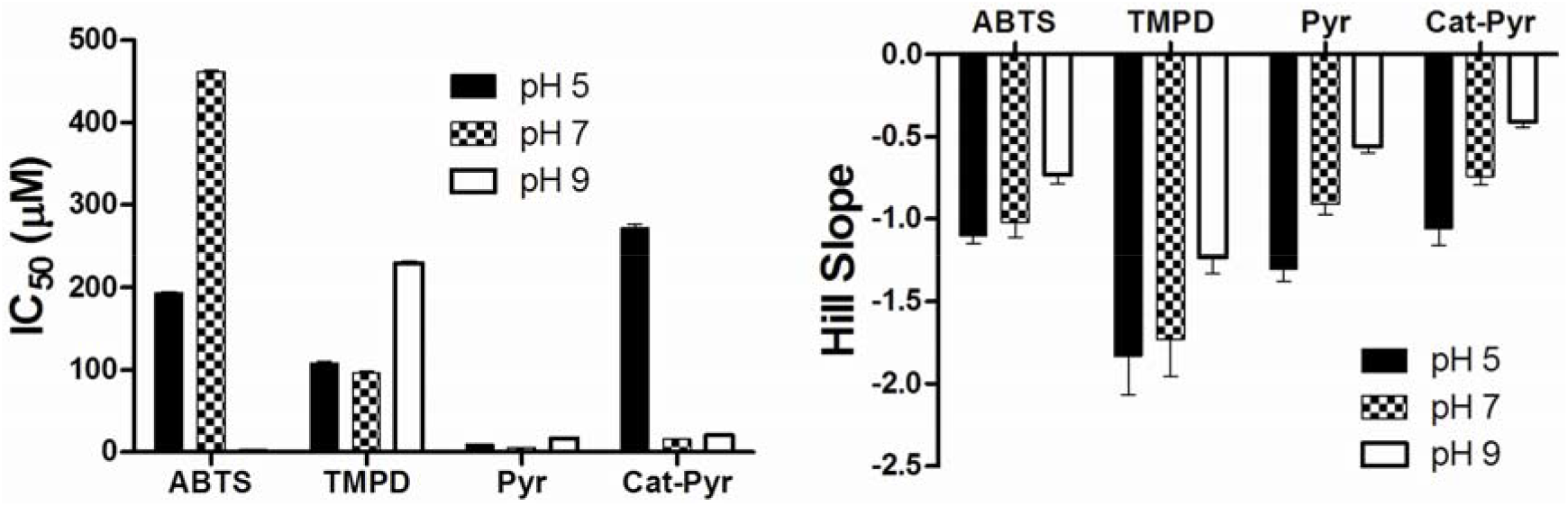
Analysis of earlier reports for the cyanide mediated IC_50_ and Hill slopes for the one-electron oxidation of diverse substrates by HRP (and Catalase – pyrogallol) at different proton availability^18,19^.

## Discussion

### The basics of the two explanations

The reaction mechanism of heme enzymes can be seen under the purview of classical (A) and murburn (B) perceptions shown in Figure 4. The classical scheme A is fastidious, requiring sequential and ordered formation of ternary complexes with high affinity binding of substrates. Post substrate binding and formation of the reactive intermediate at the heme center, the substrate MUST acquire a bonding distance with the hemeFe. Such a scheme would entail high enantioselectivity, must show exceptional substrate preferences, and would require high enzyme and substrate concentrations/ratios. These requisites are often unmet in both physiological and *in vitro* heme enzyme reactions. The murburn scheme includes the classical purview (at high enzyme substrate concentrations and ratios) and extends beyond, to low enzyme concentrations and to regimes wherein enzymes are embedded in phospholipid membranes. The interactive equilibrium espoused by the murburn scheme would also have inherent constitutive controls and also explain the atypical kinetics, diverse substrate preferences (or lack of preferences thereof!), low or lack of enantioselectivity, variable stoichiometry and explain the variability in results across labs (as minute levels of additives could significantly affect the reaction outcomes), maverick dose responses, diversity in reaction types (hydroxylation, (de)halogenation, phosphorylation, epoxidation, hetero-atom dealkylation, peroxide dismutation, etc.), and above all, account for the fast and efficient reactions at low enzyme concentrations. Unlike the classical scheme which deems DROS as an undesired toxic waste product, the murburn scheme sees DROS (particularly, the radical species) as quintessential for electron/group transfers and catalyses. Further, while the classical scheme sees water formation at heme center as the mechanism to explain for redox equivalents’ depletion, the murburn model deems stochastic DROS interactions in the milieu as the reason for redox loss/heat generation.

**Figure 4:**
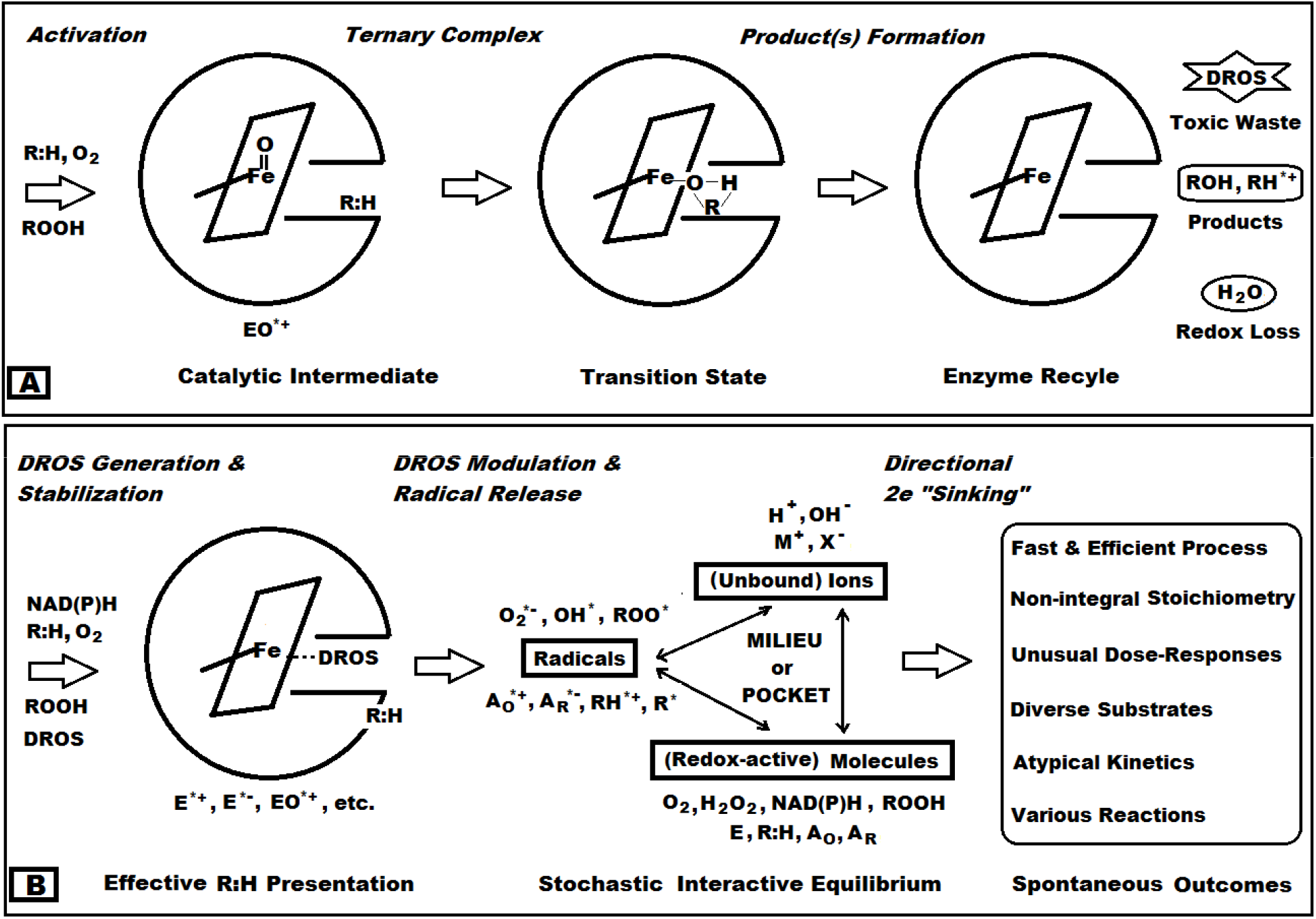
A summary of the classical (A) and murburn (B) schemes for heme-enzyme (peroxidise, P450, catalase, etc.) reactions. In A, post the charging of the heme center by the activators, the enzyme forms a 2e-deficient intermediate, which directly interacts with the final substrate bound at the distal heme pocket. DROS like superoxide and hydroxyl radicals are deemed as wasteful reaction products in this scheme. In B, the heme center serves as a DROS stabilizer cum modulator and the apoprotein would serve to enhance reaction efficiency by presenting the substrate (bound at an allosteric site, preferably adjacent to the site of DROS release. The salient feature of the murburn scheme is an interactive equilibrium between redox-active molecules. This scheme is inclusive of the classical concepts but also sees reactions outside the distal pocket as a means to affect catalysis. (A_O_ and A_R_ are oxidizable and reducible additives, M^+^ and X^−^ are positively and negatively charged ions respectively.)

Before analyzing the results of our experiments and values of K_d_-IC_50_, it is opportune to recollect the key findings from our group on diverse redox enzymes belonging to soluble peroxidases [heme histidylate species like horseradish peroxidase (HRP) and heme thiolate species like chloroperoxidase (CPO)], soluble catalases (heme tyrosylate species), and the membrane-bound mixed oxidase system of cytochrome P450s (CYPs, heme thiolate species) cum reductases (flavin enzyme).

(i) The heme distal pocket concentration plays a higher order role on the formation of a hemeFe-ligand complex. The access to hemeFe is more viable when enzyme is taken at ≥μM concentrations and at higher higher ligand:heme ratios. When the enzyme is at low concentration or embedded in membranes and when the ligand is at low concentrations, the access to ionic species and large substrates is challenged by diffusion, solubility and size constraints. For example-(a) Cumene hydroperoxide’s access to CPO’s hemeFe is limited by size constraints and therefore, this peroxide does not activate the enzyme and form Compound I. However, the very same enzyme can chlorinate molecules of much larger dimensions than cumene hydroperoxide, as the latter reaction occurs via a diffusible species. (b) HRP/CPO (10 nM) do not significantly consume 100 μM peroxide by the classical Compound I route. However, with the presence of a heme distal pocket excluded substrate like ABTS, the enzyme milieu can deplete the small peroxide activator. (This is by virtue of the “thermodynamic pull” exerted by the reactions in milieu, as espoused by murburn concept.) The formation of 2e deficient enzyme complex (the classical scheme of Compound I formation) is viable only in select scenarios (examples-*in vitro* synthetic reactions or spectroscopy sample preparations).

(ii) Ligands like CN, azide and several other molecules and ions (which are oxidizable or reducible additives) can also serve as pseudo-substrates and non-specific redox agents, helping or disrupting electron and group transfers in milieu.

Based on such findings, we have repeatedly pointed out-(a) the radical DROS’ ability to serve as electron carriers in milieu, (b) several examples of inhibitions and activations sponsored by additives as a result of events occurring beyond the hemeFe center and (c) the role of diffusible reactive species and competing reactions in milieu as the reason for varying stoichiometries and atypical kinetics in these redox enzyme reaction systems. The kinetic data and profiles obtained are therefore, more a reflection of the diffusible species’ interactive equilibriums with the enzyme/substrate/additive. It is in this context that the physiological effect of CN can be understood and explained.

Although the data points in the IC_50_ study can be plotted with acceptable R^2^ values, many heme-enzyme reactions are not actually based in the classical Enzyme-Substrate binding-based concept.^18,19^ This is evident in several profiles where data points either fall significantly below or rise above the “expected fit”, as also noted in the current work (Figure 2). Also, Ockham’s razor argues against the supposition that all such diverse enzymes have multiple modalities of binding/interactions with the “unknown and diverse substrates” and CN at the very hemeFe center or distal pocket. Further, decreasing the pyrogallol concentration lowered the IC_50_ value of cyanide for the HRP reaction at pH 5, confirming that the effect is NOT due to CN and pyrogallol competitively interacting at the distal heme active site pocket.^18^ Furthermore, we have also demonstrated with HRP-TMPD reactions that even with drastically substoichiometric hemeFe: CN ratio of 5 nM: 1.0 to 0.01 nM, significant inhibition is derived.^18^ Finally, analyses of data obtained from several heme enzymes + additives systems lead to a simple deduction-there exist multiple competing reactions and interactive redox equilibriums in the milieu, and protons are a highly crucial reactant/catalyst.^9,10,18,19^ Else, we cannot reason observations such as-IC_50_ for CPO-TMPD reaction is practically insensitive to decades of mM CN concentration as the reaction pH approaches TMPD’s pK_a_ value.^18,19^ Since radicals are stabilized at low concentrations and a radical like superoxide is more stable at low proton levels, the enhanced inhibitory ability at low cyanide concentration at pH 9 is explained. In Figure 2, the abrupt fall in activity levels of catalase/peroxidase from 1 mM to 3 mM CN is owing to the enhancement of murburn scheme by factual heme-binding of cyanide (thereby enhancing the availability of cyanide near the vicinity of heme), making it an even more potent inhibitor.

In the classical purview, CN can potentially interact with the hemeFe center via the distal heme pocket prior/post the activation step (thereby, competing with the initial activator or final substrate) and also interact with an allosteric site on the protein (bringing about an influence through non-competitive mode). The latter non-competitive modality and the uncompetitive mechanism (CN binding only to the ES complex) are non-defensible from a theoretical/practical perspective. (The experimental binding is seen at the heme-center, with only the free enzyme! So, the competitive inhibition MUST result, if the binding based explanation was true.) However, quite like Cox, even the model enzymes employed here-peroxidase (HRP) and catalase, are known to show a “non-competitive” inhibition outcome with cyanide. Therefore, the binding-based explanation stands falsified by “proof of contradiction” (along the lines employed in the K_M_ vs. K_d_ discrepancy obtained from experimental kinetics data for Cox) and the “multiple competitive redox reactions in milieu with proton as a crucial limiting reactant” scheme of murburn explanation is deduced as the reason for the outcomes. On the other hand, if DROS served as the catalytic agents and the reactions did not occur at the hemeFe alone (as per the more comprehensive perspective that murburn scheme espouses), then the competitive reactions occurring in milieu (wherein CN is present and could affect outcomes) or pocket would reflect in the experimentally derived kinetic data as “non-competitive” inhibition.

As reasoned from the founding experiments of murburn concept,^34^ the addition of DROS (superoxide) in a single shot at one locus (via a pipette tip) does not adequately capture the rates or selectivity of physiological activity of the redox enzymes wherein the DROS are dynamically generated over time and stabilized at discrete loci (in this context-by the proteins embedded in the inner mitochondrial membrane). In the current study, we have demonstrated inhibition by cyanide at mM levels in bulk aqueous milieu (which is nevertheless, better than the actual binding efficiency of cyanide with heme enzymes!). In actual physiology, as the mitochondrial membrane and proteins ensure a proton limitation cum CN^−^ confinement and effective ADP presentation, the inhibitory effect of CN ion-radical would be magnified by many orders. This is corroborated by the demonstration herein and elsewhere earlier that inhibition of heme-enzyme/DROS mediated one-electron oxidation of several molecules is critically dependent on proton availability in milieu.^9,18–19^

### Explaining the reaction chemistry at Complex IV and the effect of toxic gases

While the mechanism of toxicity of CO is understood to be related to decrease in energy levels and change in redox state owing to metabolic acidosis,^35^ the acutely toxic effect of small doses of CN is rather unknown. The research in this area is fraught with ethical issues and want of literature. Only conjectures exist that the CN toxicity results owing to its ability to bind Cox. Unlike CO, however, hypoxia is not the reason for the lethal toxic effects of CN.^35^

Wharton and Gibson had compared the oxidation of *Pseudomonad* Cox heme by O_2_ and CO binding processes.^21^ They found that the on-rate was 5.7 × 10^4^ M^−1^ s^−1^ for O_2_, thrice the rate for CO (in similar ranges). The off rate for O_2_ was less than four folds the same for CO (once again, in similar ranges). Other researchers also noted similar results.^21^ Therefore, unless the protein’s active site provides great distinction abilities to choose amongst the ligands, stark demarcations in binding are not expected across the various heme enzymes. Since there is little evolutionary pressure on Complex IV or hemoglobin to preferentially bind cyanide, the biological outcome observed with these ligands must be explained differently. If binding-based effects were the determinants, then the greater toxicity of CN (when compared to CO or H_2_S) cannot be explained. On the other hand, if the proteins evolved for a fast binding of oxygen/DROS and efficient utilization thereafter, the equilibrium and kinetic constants of Cox with O_2_ would make better physiological sense.

In our recent works with (sub)micromolar levels of heme-enzymes and several toxic small molecules and ions (N-heterocyclics, cyanide/azide, etc.), we had shown/argued that the latter are more likely to act as pseudo-substrates in milieu.^18,19,36–38^ This implies that only at higher heme?ligand ratios and high enzyme concentrations, an ion like cyanide serves as an efficient active-site ligand. The *in situ* or *in vitro* assays show a low functional IC_50_ or pseudo-K_i_ or pseudo-K_M_ (~10^−6^ M) for certain enzyme reactions,^18,19,36–38^ and such observations capture the essence of CN’s physiological toxicity. We had recently proposed that DROS are ultimately and obligatorily required as catalytic agents for metabolic activities and electron transfer (not just as molecular messengers); particularly for ATP-synthesis and heat generation in mitochondria.^3–411,18,36^ Given that the chemiosmotic explanation for cellular respiration is untenable, the thought lines that “the toxicity difference must primarily lie in DROS reaction/interaction dynamics” makes better rationale, particularly under the light of the structure-function correlations of mitochondrial proteins. The mitochondrial respiratory proteins (Complexes I through IV) show ADP binding sites, O_2_ accessible redox centers/channels, and the ability to produce DROS.^6^ Therefore, the relevant scheme for various protagonists’ interaction with Cox is projected in Figure 5. In the physiological schemes (at micromolar levels of cyanide), Cox (or other respiratory proteins) are rather unperturbed in their activity for directly interacting with O_2_/DROS. (It can also be seen from Table 1 that *k*_on_ is the highest for O_2_ and it must be remembered that oxygen is available at much higher concentrations and more readily than CN!)

**Figure 5:**
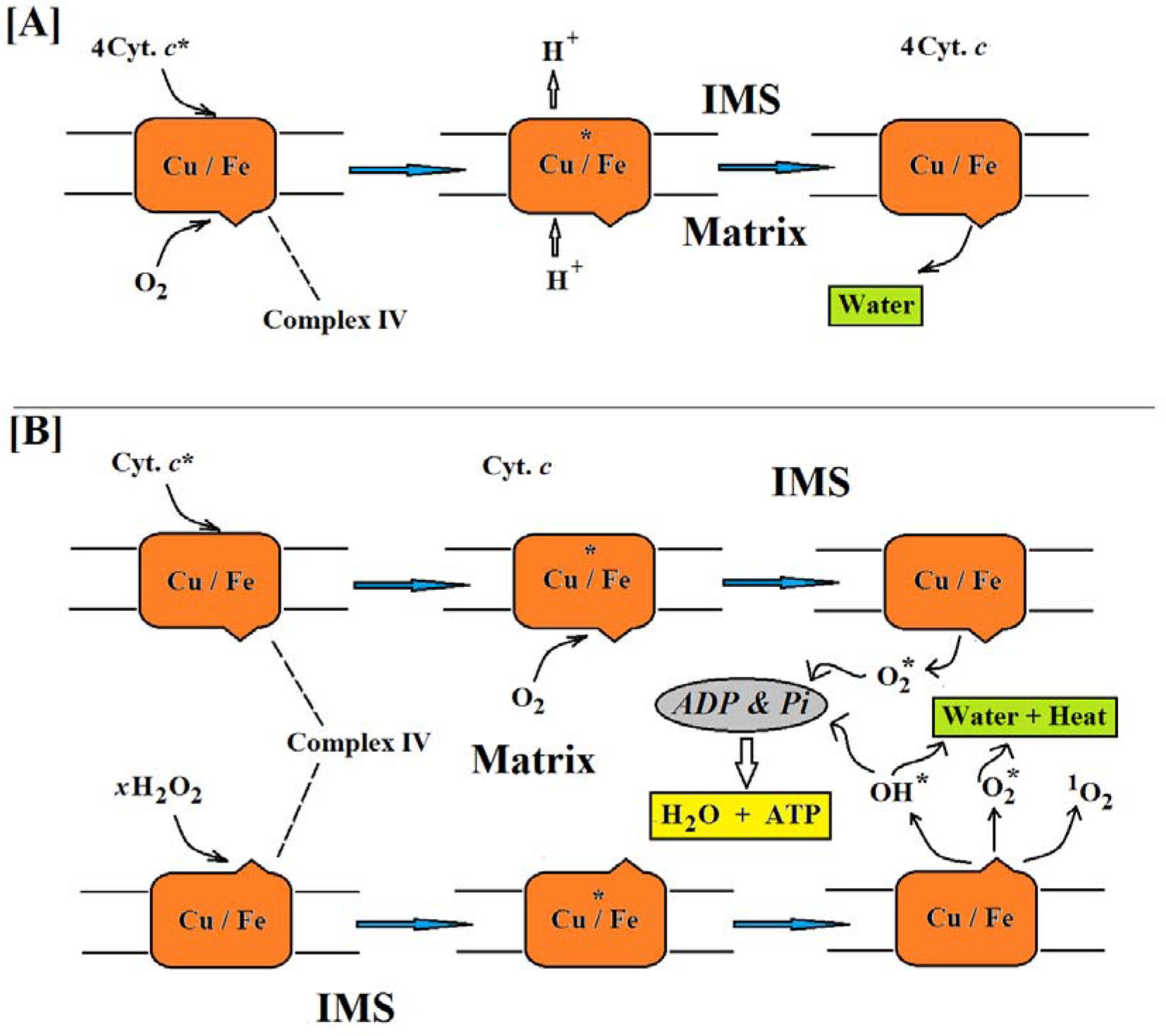
Complex IV’s proposed interaction with cytochrome c, oxygen and peroxide: **[A]** In the erstwhile mechanism, oxygen merely bound to Complex IV to get reduced by four electrons from four cytochrome c (Cyt. c) molecules and pump out protons on the way to making water at the active site. The hemeFe goes through the two-electron deficient Compound I route. **[B]** In the new scheme as shown, interaction of both Cyt. c and peroxide is to aid a one-electron scheme within the matrix. The DROS produced attack the ADP (bound on the enzyme) and Pi in milieu to make ATP. The DROS that do not get to react with ADP/Pi react among themselves to make water and heat. A small amount of water formation is possible via Complex IV, but copper may be present to prevent this and to facilitate singlet oxygen formation (which can better activate flavins). Details of stoichiometry, electron and proton involvement are not shown. [IMS: inter-membrane space]

Therefore, reiterating our earlier assertions, the equilibrium and kinetic constants determined for such “murzymes” do not have the same significance as the values interpreted from classical theories of “conventional enzymes” and “substrates” interactions. This is because the kinetic outcome is a reflection of the diffusible intermediates’ reaction with the final substrate. It is in this final step where CN comes to play, wherein it interrupts the DROS-aided synthesis of ATP within the matrix (shown in yellow box). Moreover, if the toxicity of HCN was associated with its metabolism, then we could have a suitable explanation for the dramatic effect of cyanide on the fundamental powering logic of life. We had proposed the following catalytic role (shown in Box 2) for HCN/CN^−^ equilibrium in mitochondrial toxicity:^5–6^

#### Box 2: The disruptive catalysis by CN and comparison of reaction-based inhibitory roles of toxic gases HCN and H_2_S in physiological/mitochondrial milieu.

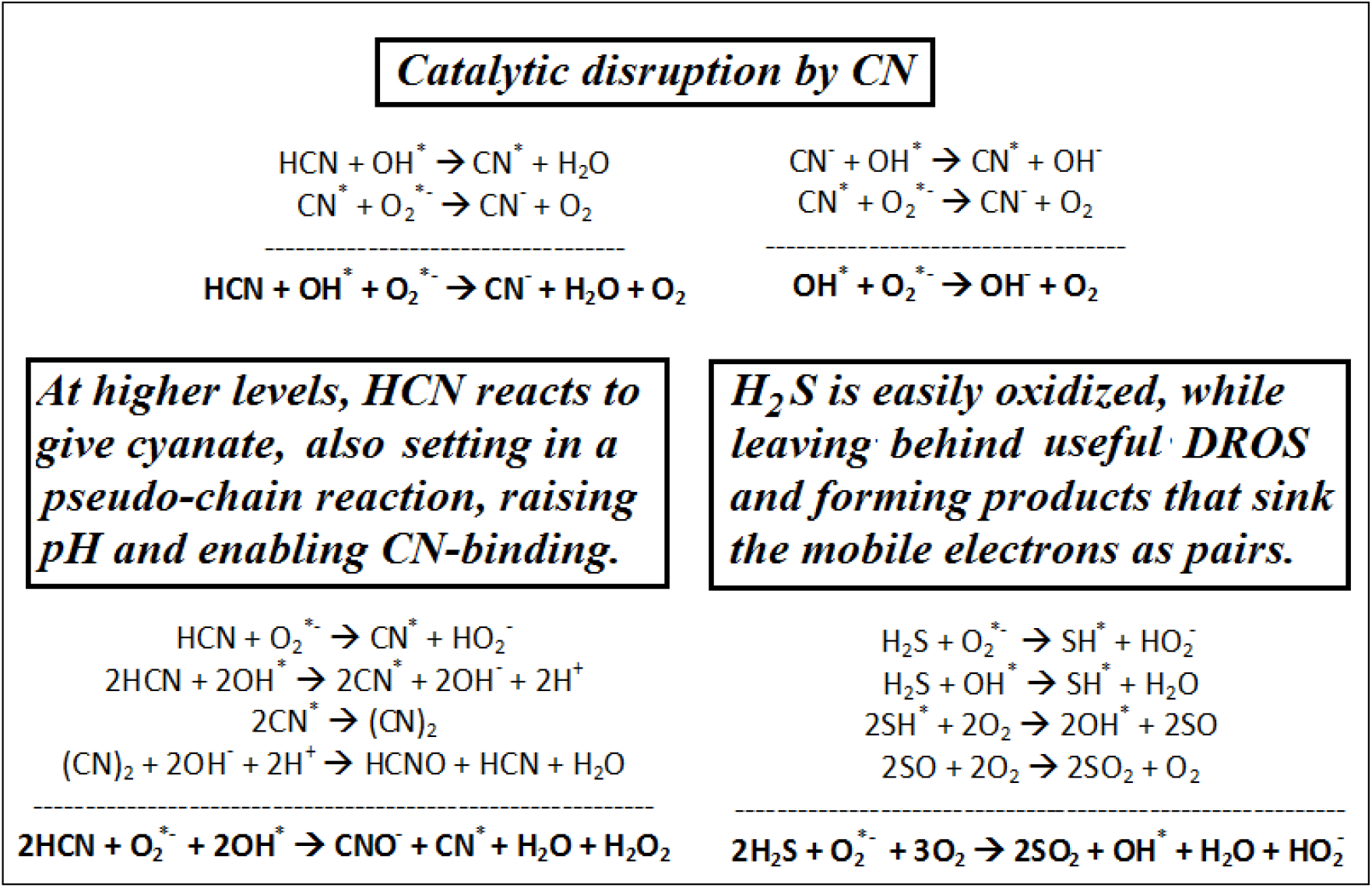

As shown, CN can effectively divert the physiological DROS utilization, forming non-productive two-electron sinks. In this murburn purview, in the steady state, protons coming in through Comp. V would be used for the neutralization of hydroxide or cyanide anions, and this outcome sabotages the cellular energy generation and homeostasis. In short, within the old binding-based purview, low amount of CN is a **SUICIDE INHIBITOR**, getting consumed in the process. In the new murburn explanation, low levels of CN (radical) can effectively serve as a **CATALYTIC INHIBITOR**, which does not get consumed. ***Further, CN is only involved with Complex IV in the erstwhile perception, whereas the murburn concept vouches that CN inhibits the ATP synthesis mediated by DROS at/around the respiratory Complexes I-IV. Therefore, CN is a global and potent inhibitor of mitochondrial ATP synthesis***.^8^ At low levels (≤ 10^−6^ M), via fast reactions occurring in the matrix, “CN molecule – CN anion – CN radical” equilibrium effectively dissipates the functional DROS. At physiological pH, HCN easily permeates phospholipid membranes and gets converted and entrapped as CN^−^ within the mitochondria (owing to highly negative charge densities in the mitochondrial membrane, afforded by cardiolipin). This enables the futile recycling of DROS, forming hydroxide, which is neutralized by the inflow of protons via Complex V. This effect scuttles the fundamental energetics’ machine logic of cells.

With latency and at higher cyanide levels (Box 2), clinical cyanosis could result. This is due to the production of the basic cyanate ion, which increases pH, enabling cyanide ion ligation to hemeFe. (Systemic lactic acidosis could further result thereafter, owing to higher order cellular machinery’s efforts to generate energy currency via anoxic modality.) Therefore, death and cyanosis owes more to the production of cyanide radical and not owing to the original binding affinity of cyanide ion or HCN, perse. In heme-enzyme systems, EPR spectroscopy has evidenced the presence of cyanide radicals and the formation of cyanate (from cyanide) has also been reported.^39,40^ Further, the lower toxicity of H_2_S is explained by the new proposal because of the facile oxidation of H_2_S to SO_2_ (which could be further oxidized to sulphate, as sulfur moieties are efficient nucleophiles) via a radical process that still leaves catalytically useful DROS in milieu (Box 2).

The new proposal is supported by the fact that HCN and H_2_S are metabolized in animal and plant systems.^41^ The lethal dose and acute toxicity predicted by the murburn chemistry scheme would be CN > CO > H_2_S, which is in fact the physiological case. Very importantly, the biggest evidence favouring the murburn explanation of CN toxicity is that unlike the competitive inhibitor of CO (which is much less toxic!), CN and H_2_S are seen/known as non-competitive inhibitors^17^. Also, it cannot be coincidental that CN is seen to inhibit the oxidase activity of model systems of catalase-peroxidase via a non-competitive purview.^42,43^ The K_d_ values of HCN and H_2_S for Cox are much higher than their Ki, which is an inexplicable theoretical premise. Further, non-competitive inhibition is one in which the inhibitor is supposed to bind at an alternative site, other than the substrate. In the current scenario, there is little scope to envisage for an “allosteric binding” based conformation or mechanistic change. Such an evolutionary pressure does not exist. This is yet another solid support for the DROS-based metabolic reaction mechanism that occurs outside the active site. ***Bottom line-DROS (radical) reactions are fast, with rate constant k values of 10^8^ to 10^10^ M^−1^ s^−1^ (about six orders higher than CN’s heme-binding rates!), which could potentially explain the rapid/acute toxicity of cyanide (radical). The rate constant k_on_ for cyanide interaction with hemeproteins falls in the consensus of only 10^2^ to 10^3^ M^1^ s^−1^. Multiplying this second order constant with micromolar concentrations of CN gives a process with rates of <10^−3^ s^−1^. That is, the binding-based activity should take at least tens of minutes to hours (depending on the variables involved), to result in cell/organism death. Therefore, such a slow and reversible binding of CN cannot explain the fast/acute lethality of cyanide that results in seconds to minutes. Radical formation and their reactions in mitochondria are thermodynamically and kinetically facile.^5^ Further, the low K_M_ for O_2_ and low Ki for the inhibitors HCN and H_2_S are the result of practically diffusion limited and zeroth order reaction of DROS with the various ions and molecules (that serve as pseudo-substrates)***.

**Further predictability and verifiability of the hypotheses:** Via the papers cited, we have theoretically and experimentally consolidated the idea already that DROS are obligatorily required in cellular metabolic schemes (which transpire via a one-electron radical chemistry) and also demonstrated that DROS-modulating species can have potentially high impact in heme/flavin-enzyme systems’ reaction outcomes.^44–47^ Furthermore, several experiments were recently suggested for verifying the proposals and tracing the influence of cyanide in cellular respiration.^48^ Murburn concept predicts a “maverick” concentration dependent effect in physiological regimes (wherein DROS are liberated in a “sustained release” fashion). We have shown and also postulated that very small amounts of cyanide may in fact enhance reaction outcomes.^19,48^ A key interesting observation noted in this regard from literature is that Cayman chemicals brochure^31^ shows that nanomolar levels of cyanide enhances activity of Cox. This result could be explained by the concentration-dependent rate variability of reactions involved in the molecule-ion-radical equilibrium, as shown in Box 2.

## Conclusions

The CN-Fe binding-based treatment does not explain the dosage regimes. Also, thermodynamics and kinetics of hemeFe-CN binding-based outcomes do not account for the physiological toxicity of cyanide. Further, the comparison with H_2_S and CO leads to the inference that a more discretized and irreversible catalytic reaction logic (and not a localized/isolated reversible effect) is the likely causation for the acute toxicity of cyanide. It is now an established idea that unusual *in vitro, in situ* and physiological dose responses mediated by low concentrations of diverse additives (like redox active small molecules, vitamins, phenolics, N-heterocyclics, azide, etc.) are mediated via the formation of catalytic radicals. Quite analogously, it is proposed that acute lethality of cyanide results due to an ion-radical equilibrium/reaction chemistry. Lethality can be defined as a status where a cell “fails to commission the works needed to sustain life”. While oxygen deprivation does not lead to loss of consciousness or lethality for several minutes, administration of cyanide knocks out an animal or cell within seconds. ATP is the known chemical energy currency of the cell, and in the absence of DROS-assisted synthesis of ATP (resulting due to CN mediated catalysis), Complex V’s high ATPase activity depletes the cellular reserves of energy. As a result, active powering mechanisms to retain the cellular functions are lost with the presence of cyanide. This inference is strongly supported by CN’s ability to inhibit DROS-assisted ATP synthesis, as predicted by murburn purview^48^ and amply demonstrated in the current work. Also, the explanation for CN-binding to proteins must be a competitive inhibition process. Going by experimental data reported till date, this is clearly not the case. It is most crucial to note the fact that murburn scheme is a “non-competitive” scheme that explains the kinetics of acute cyanide toxicity. The ideas proposed herein may potentially aid cyanide-poisoning therapy and also qualify biochemical research.

## Acknowledgments

The work was powered by Satyamjayatu: The Science & Ethics Foundation and Dept, of Biotechnology, Govt, of India grant no. BT/PR16100/NER/95/73/2015. Inputs from Alexander Scheeline (UIUC, USA), Thomas Poulos (UCI, USA), Daniel Andrew Gideon (Bishop Heber College, Trichy, India) and Vivian David Jacob (Bioculer, India) are acknowledged.

## Materials and Methods

The manuscript probes the quantitative aspects (of stoichiometry, thermodynamics and kinetics) for binding-based explanation of various ligands/gases with heme proteins based on the analysis of available literature. Further, direct experimental methods are employed to show-(i) the dynamics of competitive reactions in heme-enzyme milieu and the effect of protons on the overall outcomes. & (ii) synthesis of DROS-mediated synthesis of ATP from ADP+Pi and CN’s ability to inhibit this reaction.

### Cyanide’s impact on functional K_d_ estimation in peroxidase-catalase systems

The details of enzyme assays (molar extinction coefficients, overall workflow, etc.) have been published in detail earlier elsewhere.^18,19^ HRP and bovine catalase were product numbers P-6782 and C-9322 respectively, procured from Sigma Chemicals, USA. Other chemicals were purchased from reputed providers and were of analytical grade. Reactions were carried out in disposable cuvettes, and the concentrations of components are given in the particular figure legends. Graphpad Prism 5.02 was used to plot and analyze the data for slopes and IC_50_ values.

### Superoxide mediated ATP synthesis and inhibition by CN

KO_2_ was purchased from Sigma Aldrich, Na-ATP and NaCN was from SRL Chemicals, Na-ADP and DMSO was from HiMedia, India. 20 mg KO_2_ was of dispensed into 1 ml of anhydrous DMSO just before the experiment was commenced and this solution was used as the superoxide stock. All reagents/stocks were filtered/centrifuged to remove particulate matter. Reactions of a total of 1 ml volume were carried out as per the protocol reported earlier by our group.^8^ Experiments were conducted in 2 ml amber centrifuge tubes at pH 7.8 of Tris-Cl buffer (100 mM), with 5 minutes of incubation at 27 °C, after the final addition of 10 ul of superoxide stock (or pure DMSO in the respective control) to the made-up aqueous milieu. From the reaction incubate, 10 μl sample was then taken by an auto-injector for analysis by HPLC. Reversed phase HPLC was carried out with the protocol reported by Liu et al. as the guiding principle.^49^ A Shimadzu HPLC system was employed at a constant flow rate of 1.2 ml/min (~1800 psi), with a temporally programmed K-phosphate buffer (pH 7.0, 100 mM) and acetonitrile mixure, as given in Table 3 below.

**Table 3:**
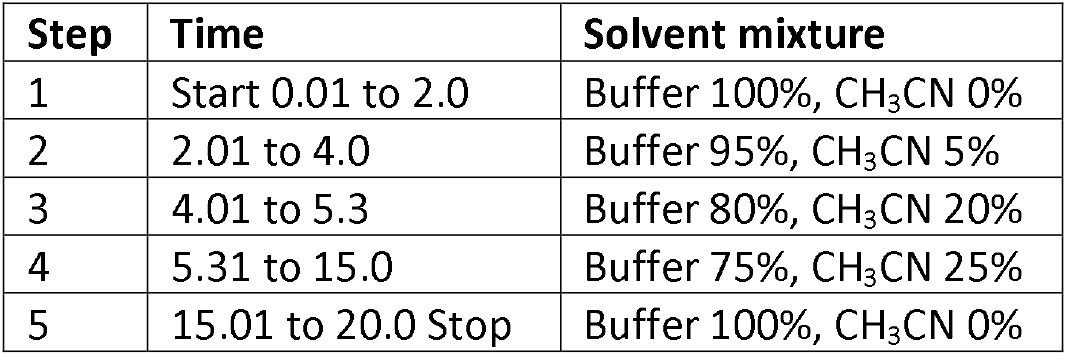
Program employed for HPLC pumps.

The reaction product was separated from the reactants using a Shim-pack GWS-C_18_ column and the chromatographic elute was analyzed online with an SPD-M30A Photodiode Array Detector at 254/220 nm for the nucleotide base signal. As shown in Figure A (top panels), under these conditions, the column voided at 2.5 to 3 minutes, whereas ATP and ADP eluted at 5.6 and 6.5 minutes respectively. The standard plot of the analytes (middle panel of Figure A) was made in the range of 0 to 200 μM. The plot gave linear regression R^2^ values >0.99; with a practically similar slope of ~6.1 × 10^6^ peak area per unit concentration (for 254 nm), for both ADP and ATP. The respective values were used for the estimation of ATP formed and ADP remaining in the reaction mix. A sample chromatogram is shown in Figure 6.

**Figure 6:**
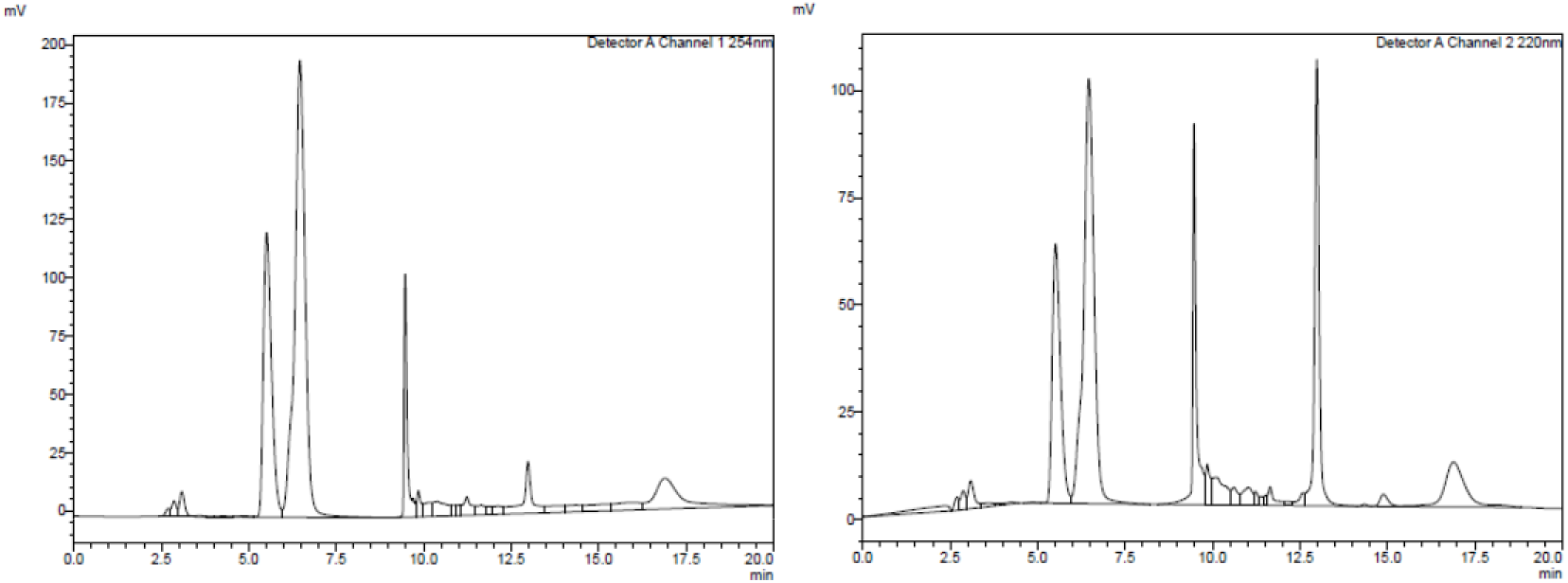
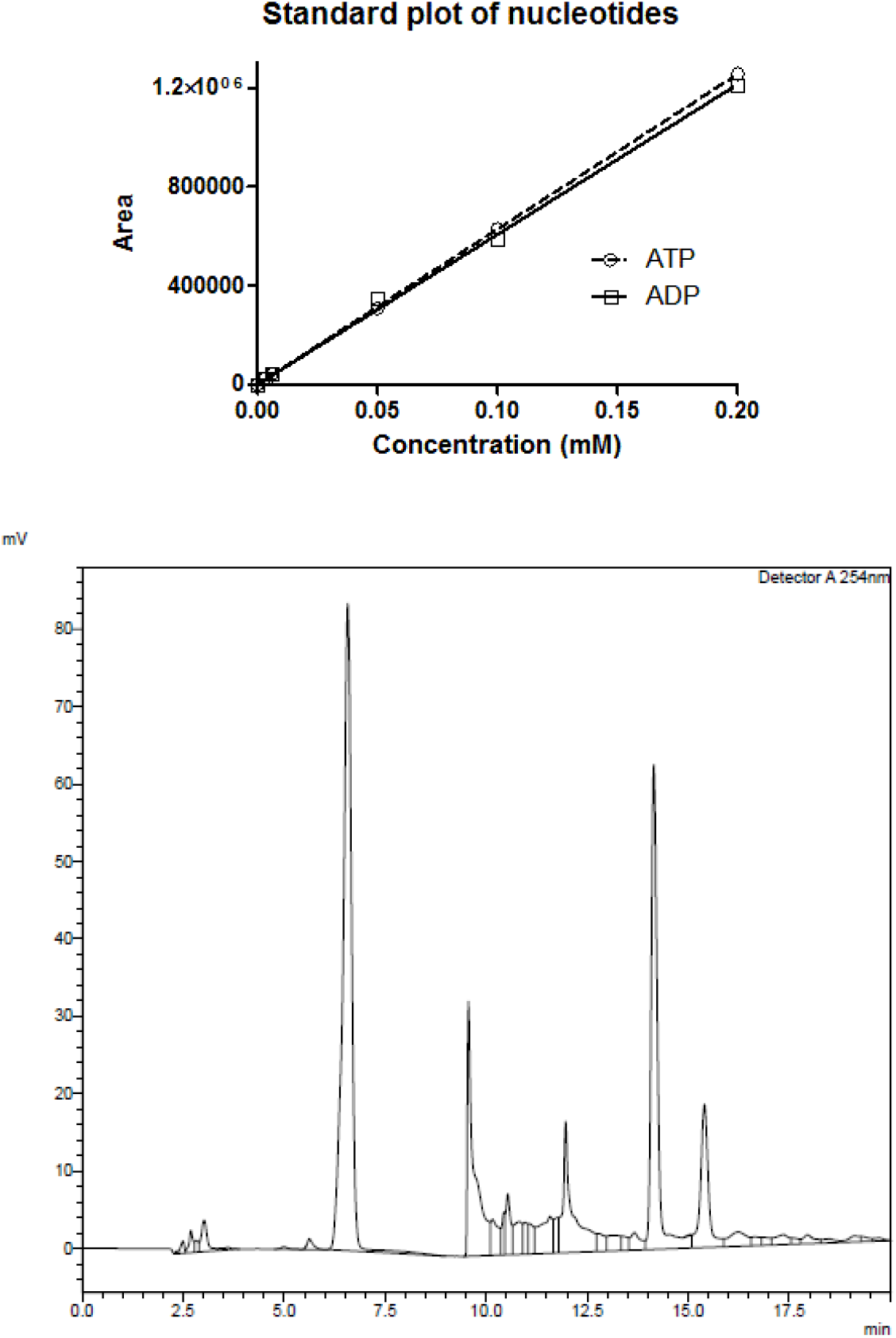
Chromatographic procedures employed for the separation, detection and estimation of ADP and ATP. The top panels show chromatographic profiles of a mixture of high amounts of ATP and ADP (0.33 and 0.76 mM respectively) at 254 nm (left) and 220 nm (right). The middle panel is the standard plot for ADP & ATP at 254 nm. The bottom panel shows a sample chromatogram (No. 4 of data Table 2).

## Supplementary Information

### Box A: The scheme for interactive equilibrium and kinetic constants for heme-ligand binding.

The superscript of * and ^±^ connote a change in charge/electronic states from the initial state. For example, heme can have Fe(ll) or Fe(lll) states and the ligand could be O_2_ or O_2_^*-^, respectively.

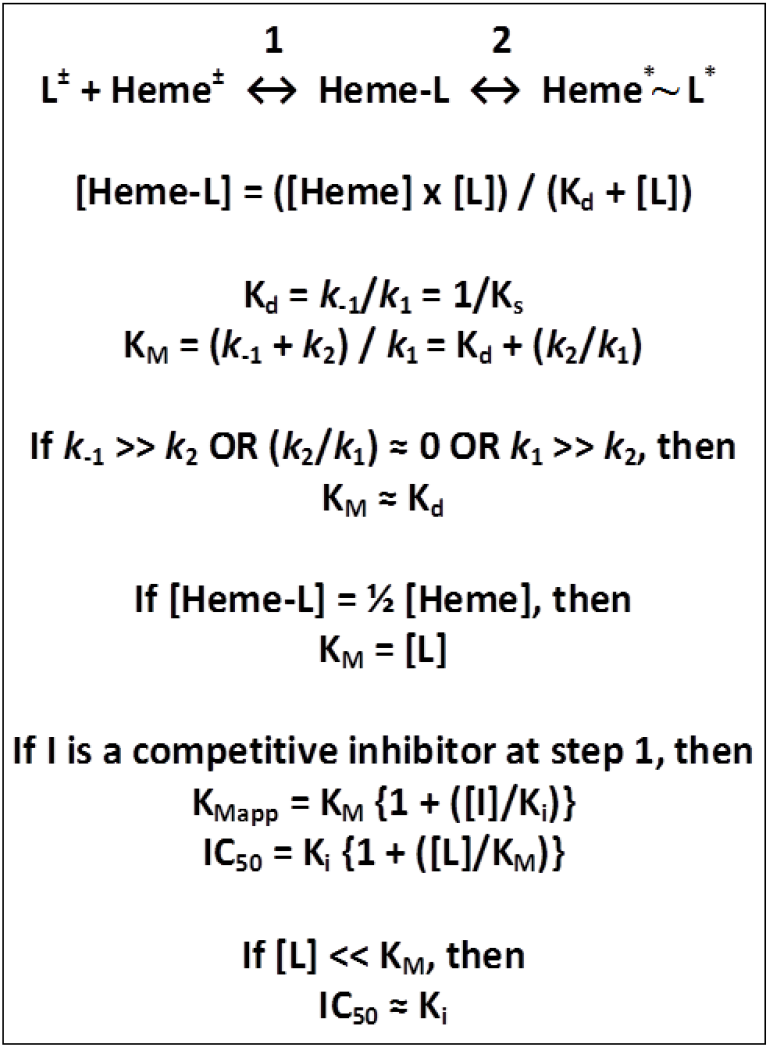

**Figure A.**
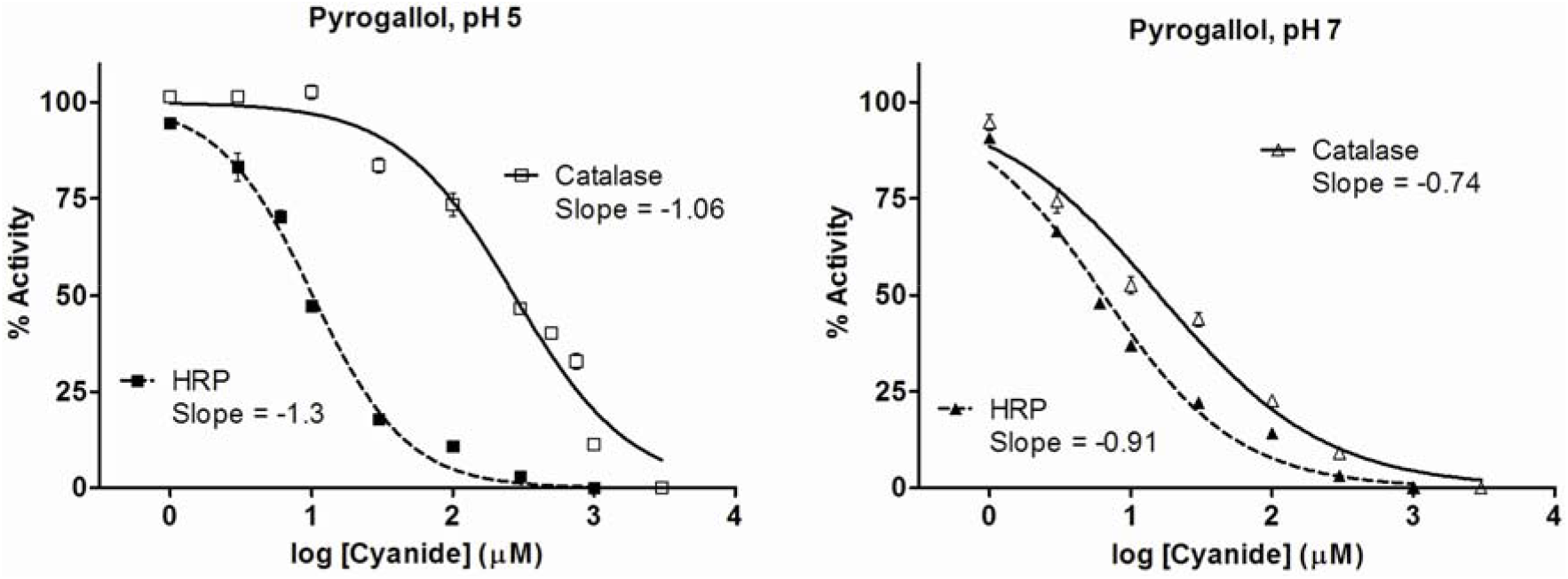
CN-inhibition profiles for Catalase/HRP mediated oxidation of pyrogallol at pH 5 and 7.^18,19^

**Table A:** Determination of functional K_d_ (extrapolation from experimental inhibition values) for data points of Figure 2 (pH 9).^18,19^. It is evident that the Kd values calculated from reaction kinetics increase upon increasing the ligand concentration, which is something that does not stand by the basic premises of the binding-based explanation.

| Total CN<br>($\mu\text{M}$) | Activity<br>(%) | Free Heme<br>(nM) | Heme-CN<br>(nM) | Free CN<br>(nM) | $K_d$<br>($\mu\text{M}$) |
| --- | --- | --- | --- | --- | --- |
| **Catalase (2 nM), pH 9** |  |  |  |  |  |
| 300 | 27 | 0.55 | 1.45 | 299999 | **113** |
| 100 | 35 | 0.69 | 1.31 | 99999 | **53** |
| 30 | 50 | 1.01 | 0.99 | 29999 | **31** |
| 10 | 61 | 1.22 | 0.78 | 9999 | **15** |
| 3 | 67 | 1.34 | 0.66 | 2999 | **6** |
| **HRP (2 nM), pH 9** |  |  |  |  |  |
| 300 | 20 | 0.40 | 1.60 | 299998 | **74** |
| 100 | 24 | 0.47 | 1.53 | 99998 | **31** |
| 30 | 45 | 0.90 | 1.10 | 29999 | **24** |
| 10 | 57 | 1.14 | 0.86 | 9999 | **13** |
| 3 | 66 | 1.32 | 0.68 | 2999 | **5.8** |
| **Catalase (2 nM), pH 7** |  |  |  |  |  |
| 100 | 22.6 | 0.45 | 1.55 | 99998.5 | **29** |
| 30 | 43.8 | 0.88 | 1.12 | 29998.9 | **23** |
| 10 | 52.6 | 1.05 | 0.95 | 9999.1 | **11** |
| 3 | 74.3 | 1.49 | 0.51 | 2999.5 | **8.7** |
| **Peroxidase (2 nM), pH 7** |  |  |  |  |  |
| 100 | 14.2 | 0.28 | 1.72 | 99998.3 | **16.6** |
| 30 | 22.2 | 0.44 | 1.56 | 29998.4 | **8.5** |
| 10 | 37.0 | 0.74 | 1.26 | 9998.7 | **5.9** |
| 3 | 48.0 | 0.96 | 1.04 | 5999.0 | **5.5** |

### Box B: Equations and schemes of some reactions/models studied herein.

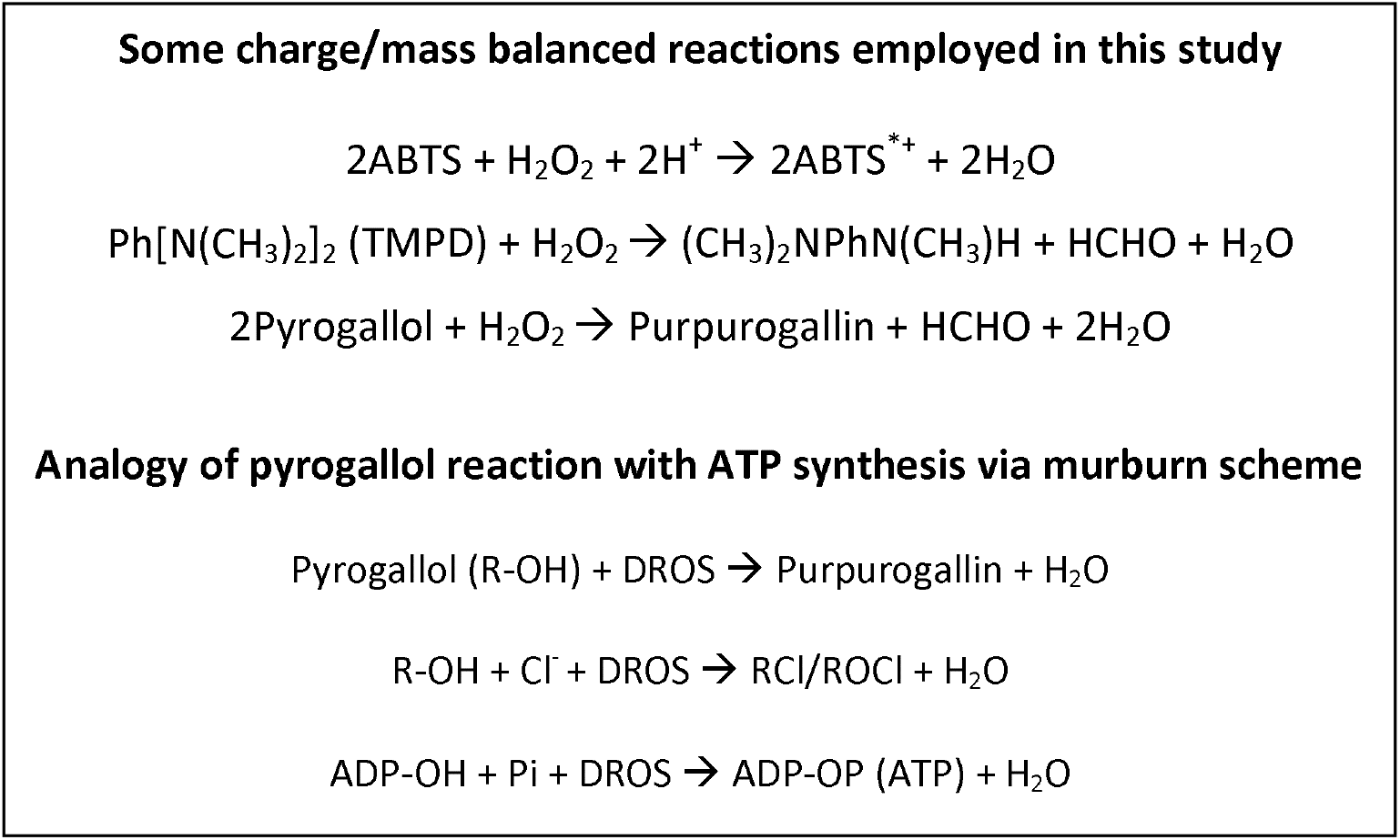

## References

1. Beasley DMG, Glass Wl. Cyanide poisoning: pathophysiology and treatment recommendations. Occup Med 1998; 48:427–431. doi:10.1093/occmed/48.7.427

2. Alberts B, Johnson A, Lewis J, Raff M, Roberts K, Walter P (1994). Molecular Biology of the Cell. New York: Garland Publishing Inc. ISBN 978-0-8153-3218-3

3. Parashar A, Gade SK, Potnuru M, Madhavan N, Manoj KM. The curious case of benzbromarone: insight into superinhibition of cytochrome P450. PLoS One 2014; 9:e89967. doi:10.1371/journal.pone.0089967

4. Venkatachalam A, Parashar A, Manoj KM. Functioning of drug-metabolizing microsomal cytochrome P450s-1. In silico probing of proteins suggest that the distal heme ‘active site’ pocket plays a relatively ‘passive role’ in some enzyme-substrate interactions. In Silico Pharmacol 2016; 4:1. doi:10.1186/s40203-016-0016-7.

5. Manoj KM, Parashar A, Gade SK, Venkatachalam A. Functioning of microsomal cytochrome P450s: Murburn concept explains the metabolism of xenobiotics in hepatocytes. Front Pharmacol 2016; 7:161. doi:10.3389/fphar.2016.00161

6. Parashar A, Gideon DA, Manoj KM. Murburn concept: A molecular explanation for hormetic and idiosyncratic dose responses. Dose Response 2018; 16:1559325818774421. doi:10.1177/1559325818774421

7. Manoj KM. Debunking chemiosmosis and proposing murburn concept as the explanation for cellular respiration. Biomed Rev 2017; 28:35–52. doi:10.14748/bmr.v28.4450

8. Manoj KM, Parashar A, Jacob VD, Ramasamy S. Aerobic Respiration: Proof of concept for the murburn perspective. J Biomol Str Dynam 2019. doi: 10.1080/07391102.2018.1552896

9. Manoj KM, Gideon DA, Jacob VD. Murburn scheme for mitochondrial thermogenesis. Biomed Rev 2018; 29, 73–82. doi: 10.14748/bmr.v29.5852

10. Manoj KM. The ubiquitous biochemical logic of murburn concept. Biomed Rev 2018; 29, 89–97. doi: 10.14748/bmr.v29.5854

11. Manoj KM. Chemiosmosis principle versus murburn concept: Why do cells need oxygen? OSF preprints 2019. doi: 10.31219/osf.io/3jq8m

12. Anseeuw K, Delvau N, Burillo-Putze G, De Iaco F, Geldner G, Holmström P. Lambert Y, Sabbe M. Cyanide poisoning by fire smoke inhalation: a European expert consensus. EurJ Emerg Med 2013; 20:2–9. doi:10.1097/mej.0b013e328357170b

13. Hall AH, Saiers J, Baud F. Which cyanide antidote? Crit Rev Toxicol 2009; 39, 541–552. doi: 10.1080/10408440802304944

14. Schwerzmann K, Cruz-Orive LM, Eggman R, Sänger A, Weibel ER. Molecular architecture of the inner membrane of mitochondria from rat liver: a combined biochemical and stereological study. J Cell Biol 1986; 102:97–103. doi:10.1083/jcb.102.1.97

15. Taylor J, Roney N, Harper C, Fransen ME, Swarts S. Toxicological profile for cyanide. Agency for Toxic Substances and Disease Registry (US) 2006. (https://www.atsdr.cdc.gov/toxprofiles/tp8.pdf)

16. Raza SK, Jaiswal DK. Mechanism of cyanide toxicity and efficacy of its antidotes. Def Sci J 1994, 41, 331–340.

17. Cooper CE, Brown GC. The inhibition of mitochondrial cytochrome oxidase by the gases carbon monoxide, nitric oxide, hydrogen cyanide and hydrogen sulfide: chemical mechanism and physiological significance. J Bioenerg Biomembr 2008; 40:533–539. doi:10.1007/s10863-008-9166-6

18. Parashar A, Venkatachalam A, Gideon DA, Manoj KM. Cyanide does more to inhibit heme enzymes, than merely serving as an active-site ligand. Biochem Biophys Res Commun 2014; 455:190–193. doi:10.1016/j.bbrc.2014.10.137

19. Manoj KM, Parashar A, Avanthika V, Goyal S, Moharana S, Singh PG, et al. Atypical profiles and modulations of heme-enzymes catalyzed outcomes by low amounts of diverse additives suggest diffusible radicals’ obligatory involvement in such redox reactions. Biochimie 2016; 125:91–111. doi:10.1016/j.biochi.2016.03.003

20. Antonini E, Brunori M, Greenwood C, Malmstrom BG, Rotilio GC. The interaction of cyanide with cytochrome oxidase. Eur J Biochem 1971; 23:396–400. doi:10.1111/j.1432-1033.1971.tb01633.x

21. Wharton DC, Gibson QH. Cytochrome oxidase from Pseudomonas aeruginosa IV. Reaction with oxygen and carbon monoxide. Biochim Biophys Acta 1976; 430: 445–453. doi:0005-2728(76)90020-7

22. Bienfait HF, Jacobs JMC, Slater EC. Mitochondrial oxygen affinity as a function of redox and phosphate potentials. Biochim Biophys Acta 1975; 376:446–457. doi:0005-2728(75)90166-8

23. Orii Y. Early molecular events in the reaction of fully reduced cytochrome oxidase with oxygen at room temperature. Chem Scr. 1988; 28A:63–69

24. Krab K, Kempe H, Wikström M. Explaining the enigmatic *K_M_* for oxygen in cytochrome c oxidase: A kinetic model. Biochim Biophys Acta 2011; 1807:348–358. 10.1016/j.bbabio.2010.12.015.

25. Manoj KM, Baburaj A, Ephraim B, Pappachan F, Maviliparambathu PP, Vijayan UK, et al. Explaining the atypical reaction profiles of heme enzymes with a novel mechanistic hypothesis and kinetic treatment. PLoS One 2010;5:e10601. doi:10.1371/journal.pone.0010601

26. Verkhovsky MI, Morgan JE, Wikstrom M. Oxygen binding and activation: Early steps in the reaction of oxygen with cytochrome c oxidase. Biochemistry 1994; 33:3079–3086. doi:10.1021/bi00176a042

27. Gibson QH, Greenwood C. Reactions of cytochrome oxidase with oxygen and carbon monoxide. Biochem J 1963; 86 541–554. doi: 10.1042/bj0860541

28. Gibson QH, Palmer G, Wharton DC. The binding of carbon monoxide by cytochrome c oxidase and the ratio of the cytochromes a and a_3_*. J Biol Chem 1965; 240:915–920

29. Marshall D, Nicholls P, Wilson M, Cooper C. A comparison of nitric oxide and hydrogen sulphide interactions with mitochondrial cytochrome c oxidase. Nitric Oxide 2012; 27:S11–S12. doi:10.1016/j.niox.2012.04.043

30. Wainio WW, Greenless J. Complexes of cytochrome c oxidase with cyanide and carbon monoxide. Arch Biochem Biophys 1960; 90:18–21. doi:10.1016/0003-9861(60)90605-6

31. Cayman Chemicals. Mitocheck^®^ Complex IV Activity assay kit. Item No. 700990, page 11. https://www.caymanchem.com/pdfs/700990.pdf

32. Li J, Noll BC, Schulz CE, Scheidt WR. Comparison of cyanide and carbon monoxide as ligands in Iron(II) porphyrinates. Angew Chem Int Ed 2009; 48-5010–5013. doi:10.1002/anie.200901434

33. National Research Council (US). *Acute exposure guideline levels for selected airborne chemicals (Volume 8)*. Carbon monoxide-acute exposure guideline levels. (Chapter 2, Table 2.3) Washington (DC) (2010): National Academies Press (US). https://www.ncbi.nlm.nih.gov/books/NBK220007/

34. Manoj KM. Chlorinations catalyzed by chloroperoxidase occur via diffusible intermediate (s) and the reaction components play multiple roles in the overall process. Biochim Biophys Acta 2006; 1764, 1325–1339.

35. National Research Council (US). *Combined exposures to hydrogen cyanide and carbon monoxide in army operations: Initial report*. Mechanisms of carbon monoxide and hydrogen cyanide toxicity. (Chapter 2). Washington (DC) (2008). National Academic Press (US). https://www.nap.edu/read/12040/chapter/4

36. Andrew D, Hager L, Manoj KM. The intriguing enhancement of chloroperoxidase mediated one-electron oxidations by azide, a known active-site ligand. Biochem Biophys Res Commun 2011; 414:646–649. doi:10.1016/j.bbrc.2011.10.128

37. Manoj KM, Gade SK, Venkatachalam A, Gideon DA. Electron transfer amongst flavo- and hemo-proteins: diffusible species effect the relay processes, not p rote in–protein binding. RSC Adv 2016;6:24121–24129. doi: 10.1039/C5RA26122H

38. Manoj KM. Aerobic respiration: Criticism of the proton-centric explanation involving rotary ATP synthesis, chemiosmosis principle, proton pumps and electron transport chain. Biochem Insights 2018. doi:10.1177/1178626418818442

39. Moreno SNJ, Stolze K, Janzen EG, Mason RP. Oxidation of cyanide to the cyanyl radical by peroxidase/H_2_O_2_ systems as determined by spin trapping. Arch Biochem Biophys 1988; 265:267–271. doi:10.1016/0003-9861(88)90127-0

40. Delporte C, Zouaoui Boudjeltia K, Furtmuller PG, Maki RA, Dieu M, Noyon C, et al. Myeloperoxidase-catalyzed oxidation of cyanide to cyanate: A potential carbamylation route involved in the formation of atherosclerotic plaques? J Biol Chem 2018; 293:6374–6386. doi:10.1074/jbc.M117.801076

41. Rubright SM. Cyanide and hydrogen sulfide: A review of two blood gases, their environmental sources, and potential risks. Thesis submitted to the University of Pittsburgh (2016). http://d-scholarship.pitt.edu/30128/1/Rubright_ETD_12_2016.pdf

42. Chance B. (1952) Oxidase and peroxidase reactions in presence of dihydroxymaleic acid. J Biol Chem 1952; 197, 577–589.

43. Chance B (1949) The reaction of catalase and cyanide. J Biol Chem 1949; 179, 1299–1309.

44. Manoj KM, Gade SK & Mathew L. Cytochrome P450 reductase: a harbinger of diffusible reduced oxygen species. PLoS One 2010; 5: e13272. doi:10.1371/journal.pone.0013272

45. Parashar A, Manoj KM. Traces of certain drug molecules can enhance heme-enzyme catalytic outcomes. Biochem Biophys Res Commun 2012;417:1041–1045. doi:10.1016/j.bbrc.2011.12.090

46. Gade SK, Bhattacharya S, Manoj KM. Redox active molecules cytochrome c and vitamin C enhance heme-enzyme peroxidations by serving as non-specific agents for redox relay. Biochem Biophys Res Commun 2012;419:211–214. doi:10.1016/j.bbrc.2012.01.149

47. Gideon DA, Kumari R, Lynn AM & Manoj KM. What is the Functional Role of N-terminal Transmembrane Helices in the Metabolism Mediated by Liver Microsomal Cytochrome P450 and its Reductase? Cell Biochem Biophys 2012; 63: 35–45. doi: 10.1007/s12013-012-9339-0

48. Manoj KM, Soman V, Jacob VD, Parashar A, Gideon DA, Kumar M, Manekkathodi A, Ramasamy S, Pakshirajan K. Chemiosmosis principle versus murburn concept: *Why do cells need oxygen?* Deducing the underpinnings of aerobic respiration by mechanistic predictability. OSF Preprints, doi: 10.31219/osf.io/3jq8m

49. Liu H, Jiang YM, Luo YB, Jiang WB. A simple and rapid determination of ATP, ADP and AMP concentrations in pericarp tissue of litchi fruit by high performance liquid chromatography. Food Technol Biotechnol 2006, 44:531–534.

